# Model-Dependent Renal Phenotypes in Diabetic Kidney Disease: Comparative Histopathological Characterization of Commonly Used Animal Models

**DOI:** 10.64898/2026.07.02.736132

**Authors:** Raziyeh Rezaei, Azar Naimi, Yousof Gheisari, Zahra Ramazani, Issa Sulaiman Al-Amri, Hoda Doustmohammadi, Fatemeh Jamshidi-adegani, Sulaiman Al-Hashmi

**Affiliations:** Laboratory for Stem Cell and Regenerative Medicine, Natural and Medical Sciences Research Center, University of Nizwa, Nizwa, Oman; Regenerative Medicine Research Center, Isfahan University of Medical Sciences, Isfahan, Iran; Reproductive Sciences and Sexual Health Research Center, Department of Pathology, Isfahan University of Medical Sciences, Isfahan, Iran; Department of Biological Sciences and Chemistry, College of Arts and Sciences, University of Nizwa, Nizwa, Oman

**Keywords:** Diabetic Kidney Disease, Histopathology, Glomerulomegaly, Tubular vacuolization, Mesangial hypercellularity, and Arteriolar hyalinosis

## Abstract

**Background:** Diabetic kidney disease (DKD) remains a leading cause of end-stage renal disease worldwide, characterized by progressive structural and metabolic alterations secondary to chronic hyperglycemia. While numerous type 1 and type 2 rodent models have been developed to study the pathophysiology of DKD, no single model perfectly recapitulates the full clinical spectrum of human disease. The selection of an optimal model depends deeply on the specific research objective, as phenotypic expression and histopathological severity vary significantly across different strains and induction methods. The present study provides a comparative analysis of the renal histological of three widely utilized murine models: the chemically induced streptozotocin (STZ) model and the genetic Akita (type 1) and db/db (type 2) models.

**Methods:** Male STZ-induced (28 weeks post-induction), heterozygous Akita (28 weeks old), and db/db mice at two different age intervals (18–21 and 16–24 weeks old) were assessed. Renal injury was quantified using four light-microscopic parameters: glomerulomegaly, mesangial hypercellularity, tubular vacuolization and arteriolar hyalinosis. Due to observed discrepancies between metabolic and structural findings in the db/db strain, transmission electron microscopy (TEM) was employed for subcellular characterization.

**Results:** All models exhibited significant hyperglycemia and albuminuria. At the light-microscopic level, STZ and Akita mice demonstrated consistent and pronounced renal lesions. In contrast, db/db mice despite increasing albuminuria and obesity, light microscopy revealed heterogeneous and inconsistent histopathological changes. However, TEM analysis of db/db mice kidneys successfully captured early ultrastructural injury, including irregular glomerular basement membrane (GBM) thickening and focal podocyte foot process effacement, which were undetectable by light microscopy.

**Conclusions:** Our findings indicate that the Akita and STZ-induced models exhibit prominent structural alterations detectable by conventional light microscopy, whereas the db/db model requires ultrastructural evaluation by TEM to reliably confirm renal injury. This study underscores the limitation of routine histology in certain type 2 diabetes models and highlights the complementary value of TEM for accurate histopathological characterization. Collectively, the alternative histopathological markers identified herein offer sensitive and readily accessible indices for monitoring early-to-moderate DKD progression, providing a more robust framework for preclinical model selection and therapeutic evaluation in future studies.

## Introduction

Diabetic kidney disease (DKD), a major microvascular complication of diabetes mellitus, remains one of the leading causes of chronic kidney disease (CKD) worldwide [1]. It is defined by a progressive decline in renal function accompanied by distinct structural and histopathological alterations, including glomerular hypertrophy, mesangial expansion, glomerulosclerosis, and tubulointerstitial fibrosis [2]. While clinical investigations have substantially advanced the understanding of disease progression and therapeutic responses, limitations such as interpatient variability, prolonged disease course, and limited accessibility to renal biopsy specimens underscore the critical importance of experimental animal models in CKD research [3, 4].

Among the currently available models, streptozotocin (STZ)-induced diabetes, Akita mice, and db/db mice are widely utilized to mimic different aspects of type 1 and type 2 diabetes-associated renal injury. However, these models differ considerably in their ability to recapitulate the complex pathophysiology and histopathological spectrum of human DKD [5]. STZ is a pancreatic β-cell cytotoxic agent that induces insulin deficiency and sustained hyperglycemia through β-cell destruction mediated by DNA alkylation and oxidative stress [6]. This model typically exhibit modest albuminuria and mild to moderate renal alterations, including glomerular hypertrophy, glomerular basement membrane thickening, mesangial matrix expansion [7]. However, advanced pathological features, including pronounced glomerulosclerosis, tubular atrophy, and interstitial fibrosis, are generally limited or absent in the STZ model unless additional interventions, such as uninephrectomy, high-fat diet feeding, or genetic modifications, are incorporated to accelerate disease progression [8]. The Akita mouse is a spontaneous genetic model of type 1 diabetes caused by a dominant mutation in the insulin 2 gene (*Ins2*^*Akita*^), resulting in insulin misfolding, endoplasmic reticulum stress, β-cell apoptosis, and chronic hyperglycemia [9]. Compared with STZ-induced models, Akita mice exhibit a more stable and reproducible diabetic phenotype and develop progressive renal injury characterized by mesangial sclerosis, podocyte loss, glomerular hypertrophy, and marked albuminuria. Nevertheless, the severity of nephropathy is strongly influenced by the mouse strain background [7]. In contrast, the db/db mouse is a genetic model of type 2 diabetes harboring a mutation in the leptin receptor gene (*Lepr*^*db*^), which disrupts leptin signaling and results in hyperphagia, obesity, insulin resistance, hyperinsulinemia, and persistent hyperglycemia. This model develops early glomerular lesions, mesangial expansion, and variable degrees of tubulointerstitial injury [7]. However, similar to other diabetic models, it incompletely reproduces advanced renal fibrosis and progressive renal failure without the incorporation of additional pathogenic stressors such as hypertension induction or uninephrectomy [10].

Although numerous studies have described the classical histopathological features of DKD, such as tubulointerstitial fibrosis and glomerulosclerosis, significant challenges remain. Conventional markers still show inconsistent detectability, and poor reproducibility across different rodent models [7, 11]. Notably, the severity of advanced lesions varies considerably among experimental models when assessed by routine hematoxylin and eosin (H&E) staining under light microscopy, particularly in the early-to-moderate stages of diabetes [11]. This creates a significant gap in which severe biochemical alterations, such as marked hyperglycemia and albuminuria, often do not correspond to visible structural alterations on standard light microscopy. Consequently, there is an unmet need to identify and validate alternative histopathological markers that are easily detectable yet frequently overlooked under conventional light microscopy. These markers would substantially improve the characterization and comparison of baseline kidney injury across widely used rodent models of DKD.

To address this gap, the present study aimed to comparatively evaluate the STZ-induced model and the genetic Akita and db/db mouse models by focusing on specific histopathological parameters: glomerulomegaly, mesangial hypercellularity, tubular vacuolization, and arteriolar hyalinosis. Importantly, because conventional H&E staining under light microscopy is insufficient to detect structural alterations, transmission electron microscopy (TEM) was employed as a crucial complementary tool. By evaluating these distinct parameters and employing transmission electron microscopy to overcome the limitations of light microscopy, this study seeks to provide a more reliable, refined framework for assessing experimental DKD progression and model fidelity..

## 2. Method

### 2.1 Animal models

In this study, three murine models of diabetes mellitus were evaluated and compared, including streptozotocin (STZ)-induced diabetic mice, Akita mice, and db/db mice. All animals were housed under a 12-hour light/dark cycle with free access to food and water.

Non-fasting blood glucose levels and body weight were measured using a glucometer and precision balance, respectively, prior to anesthesia. Blood samples were obtained via tail vein sampling. These measurements were used to confirm diabetic status and to monitor metabolic changes across all experimental groups.

All animal experiments were conducted in accordance with the NIH Guide for the Care and Use of Laboratory Animals and approved by the relevant institutional ethics committees prior to study initiation (University of Nizwa Ethics Committee: VCGSR, AREC/01/202 for Akita and db/db models; Isfahan University of Medical Sciences Ethics Committee: IR.MUI.AEC.1401.046 for STZ-induced models)[12].

#### 2.1.1. Streptozotocin (STZ)-induced diabetic mice C57Bl6/j (Type 1 diabetes)

Male C57BL/6J were supplied from Royan Institute of Iran. Ten mice (8–10 weeks old) received intraperitoneal injections of streptozotocin (50 mg/kg) dissolved in sodium citrate buffer for five consecutive days, following the Animal Models of Diabetic Complications Consortium (AMDCC) protocol. To reduce hypoglycaemia, mice were provided with 10% sucrose water after each injection. three weeks after induction, non-fasting blood glucose and body weight were measured. Mice with blood glucose level exceeding 300 mg/dL were considered diabetic (n=8), while age-matched controls (n=5) were maintained under identical conditions. Animal were followed for 28 weeks to allow the development of renal pathology [13].

#### 2.1.2. Akita mice (C57BL/6-*Ins2*^*Akita*^/J, Type 1 diabetes)

Heterozygous C57BL/6-*Ins2*^*Akita*^/J mice were obtained from The Jackson Laboratory and bred with C57BL/6J females. Male offspring exhibiting hyperglycaemia after 4 weeks of age were classified as diabetic (n=6), while normoglycemic littermates served as wild-type (n=6) [14]. All animals were maintained for 28 weeks to evaluate diabetes-associated renal structural changes.

#### 2.1.3. db/db mice (BKS.Cg-*Dock7*^*m*^+/+*Lepr*^*db*^J, Type 2 diabetes)

BKS.Cg-*Dock7*^*m*^+/+*Lepr*^*db*^J mice were obtained from The Jackson Laboratory. Heterozygous breeding pairs were used to generate homozygous diabetic (db/db) and control (db/m) mice. Male mice were studied in two age groups: early-stage (18–21 weeks; n=6 db/db, n=5 db/m) and late-stage (16–24 weeks; n=7 db/db, n=4 db/m) to assess age-dependent renal pathology.

### 2.2. Urinary Protein and Albumin Measurements

Twenty-four-hour urine samples were collected using metabolic cages prior to sacrifice. Samples were centrifuged at 1500 rpm, aliquoted, and stored at -80°C for subsequent analysis. Total urinary protein was measured using the Bradford assay [15], and urinary albumin was quantified using a commercially available ELISA kit (Albuwell M, Ethos Biosciences; and ab108792, Abcam), according to the manufacturers’ instructions. Albuminuria and proteinuria levels in Akita and STZ-induced mice were normalized to body weight. Due to marked obesity in db/db mice, normalization was not applied in this group.

### 2.3. Renal Tissue Collection and Histopathological Evaluation

Mice were anesthetized with ketamine (100 mg/ kg) and xylazine (10 mg/ kg) and subsequently sacrificed. Kidneys were harvested, and the right kidney was fixed in 10% (v/v) formaldehyde, processed, embedded in paraffin, sectioned, and stained with haematoxylin and eosin (H&E) for histological analysis.

Renal histopathological changes were assessed based on predefined parameters. Initial histopathological evaluation of parameters was performed in a non-blinded manner to identify consistent pathological features in type 1 diabetic mice (STZ and Akita). Subsequently, all slides were re-coded and analysed in a blinded fashion. The pathologist remained unaware of the experimental grouping (diabetic vs. control), although model classification (STZ, Akita or db/db) was retained. Identical scoring criteria were applied throughout all analyses.

### 2.4. Transmission Electron Microscopy (TEM)

Formalin-fixed paraffin-embedded (FFPE) kidney tissues were processed for TEM analysis. Samples were deparaffinized in xylene (3 changes) and rehydrated through graded ethanol series (100%–50%). Tissues were incubated overnight in distilled water, washed in sodium cacodylate buffer (3×, 4°C), and fixed in Karnovsky’s fixative for 4 hours at 4°C.

Post-fixation was performed using 1% osmium tetroxide for 1 hour, followed by dehydration in graded acetone. Samples were infiltrated with epoxy resin (Agar 100 system), embedded in Beam capsules, and polymerized at 60–65°C for 16 hours. Semi-thin sections (0.5 µm) were stained with toluidine blue for region selection. Ultrathin sections (60–90 nm) were stained with uranyl acetate and lead citrate prior to TEM examination.

### 2.5. Statistical Analysis

Statistical analyses and data visualizations were performed using GraphPad Prism (version 10.4.1). For histopathological parameters evaluated on a binary presence/absence system, relative injury frequencies were calculated as percentages. Fisher’s exact test was utilized to analyse the proportional distribution of individual lesions between the experimental groups. The non-parametric Mann-Whitney U test was used to determine differences in total injury scores across cohorts. This test was also applied to analyze statistical differences in biochemical parameters between diabetic groups and their controls. For all analyses, statistical significance was stringently defined as a P-value < 0.05.

## 3. Result

### 3.1. Validation of DKD Establishment Across Animal Models

DKD was successfully established and validated across three animal models, as evidenced by sustained hyperglycemia, proteinuria, and albuminuria at the designated endpoints. Two distinct animal models are employed to evaluate the histopathological alterations in diabetic type 1. In the chemical type 1 diabetes model, eight out of ten mice developed sustained hyperglycemia within two weeks following STZ administration, with five untreated mice serving as normal controls. At 28 weeks post-injection, diabetic mice showed significantly elevated blood glucose levels and reduced body weight compared with controls (Fig. 1a–b). No statistically significant difference was observed in the kidney-to-body weight ratio between the STZ and control groups (Fig. 1c). Renal impairment was confirmed via urinary analysis, which demonstrated a marked, approximately 9-fold increase in 24-hour albumin excretion (Fig. 1e). Urinary total protein levels were also elevated in the STZ cohort, though this alteration did not reach the threshold for statistical significance relative to controls (Fig. 1d). Similarly, at 28 weeks of age, the genetic type 1 heterozygous Akita mice displayed severe hyperglycemia and substantial body weight loss compared with their wild-type (Fig. 1f–g). In contrast to the STZ model, diabetic Akita mice exhibited a significantly increased kidney-to-body weight ratio (Fig. 1h). These structural changes were accompanied by marked functional impairment, including an approximately 6-fold elevation in proteinuria and a greater than 10-fold increase in albuminuria (Fig. 1i–j).

**Figure 1:**
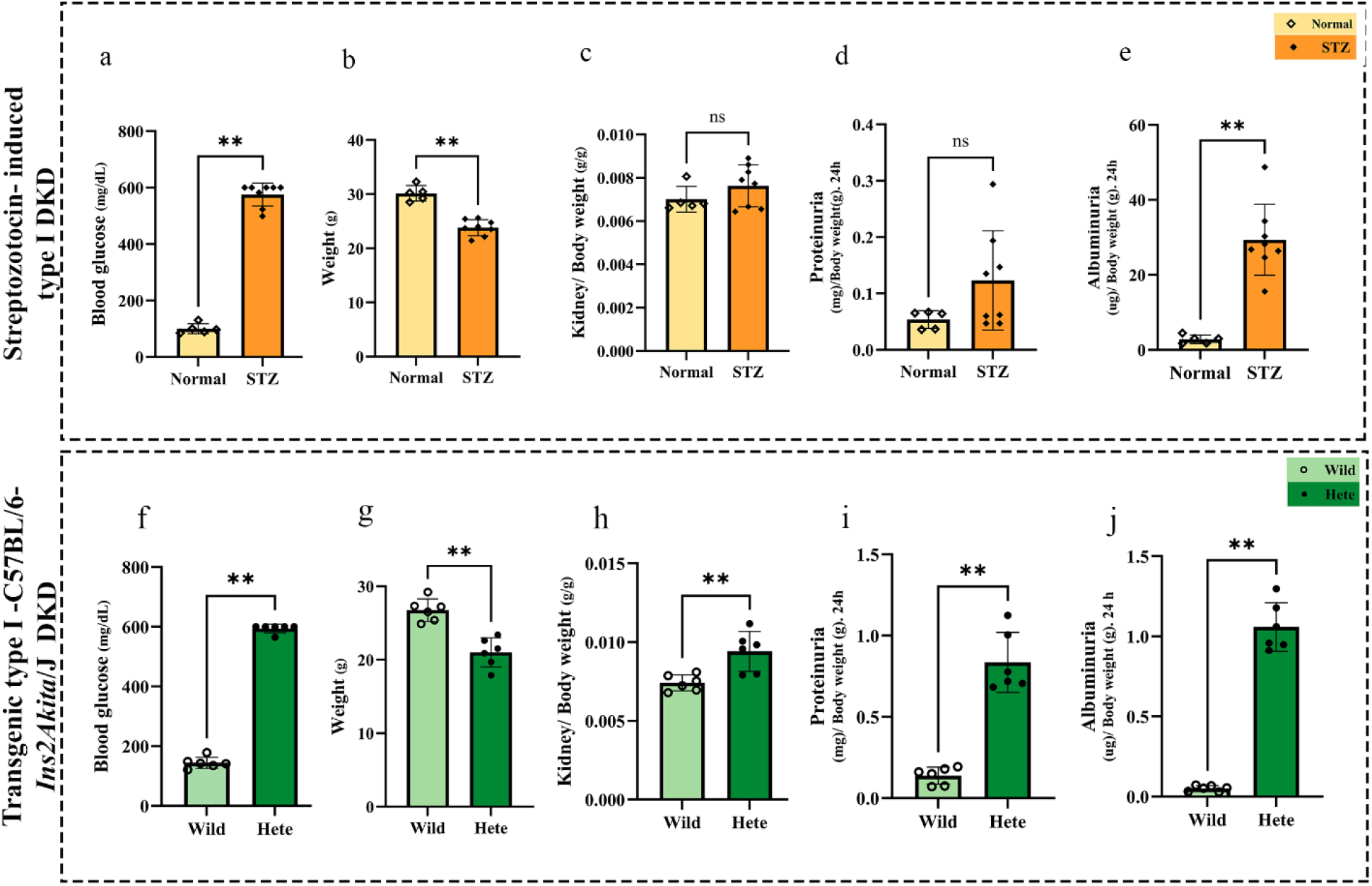
Biochemical and physiological characterization of type 1 diabetic kidney disease (DKD) animal models. Parameters evaluated 28 weeks post-induction in the STZ-induced model (n = 8 diabetic, n = 5 normal) and genetic Akita model (n = 6 per group). (a, f) non-fasting blood glucose levels, showing sustained hyperglycemia in both diabetic cohorts. (b, g) Total body weight alterations, demonstrating significant weight loss in diabetic mice compared to their respective controls. (c, h) Kidney-to-body weight ratios, revealing a significant increase uniquely in the Akita model. (d, i) 24-hour urinary total protein levels, showing a statistically significant elevation in Akita mice, while the increase in the STZ group did not reach statistical significance (ns). (e, j) 24-hour urinary albumin excretion, showing an approximate 9-fold increase in STZ and a greater than 10-fold elevation in Akita mice. Data are presented as median with range. Albuminuria was quantified via ELISA, and total protein was determined using the Bradford assay. (ns = non-significant, **P < 0.01)

For the genetic type 2 diabetes model, cohorts were categorized into early-stage (18–21 weeks old; n=6 db/db, n=5 db/m) and late-stage (16–24 weeks old; n=7 db/db, n=4 db/m) disease progression. This classification was established to capture potential stage-dependent progression, as the latter cohort exhibited significantly more advanced biochemical markers of renal injury despite the overlapping chronological age. In both age groups, db/db mice manifested severe obesity and marked hyperglycemia, showing highly significant increases in body weight and blood glucose levels relative to their lean db/m controls (Fig. 2a–b; Fig. 2f–g). The kidney-to-body weight ratio did not differ significantly between the groups in either cohort when normalized to body weight (Fig. 2c; Fig. 2h). The early-stage db/db mice demonstrated a non-significant rising trend in proteinuria (Fig. 2d), whereas the late-stage cohort developed a highly prominent and statistically significant elevation in total protein excretion (Fig. 2i). Furthermore, urinary albumin excretion was severely elevated across both cohorts, demonstrating an approximate 10-fold increase in the early-stage group and a progressive and markedly greater elevation (>20-fold) in the late-stage cohort (Fig. 2e; Fig. 2j), confirming robust progression of DKD.

**Figure 2:**
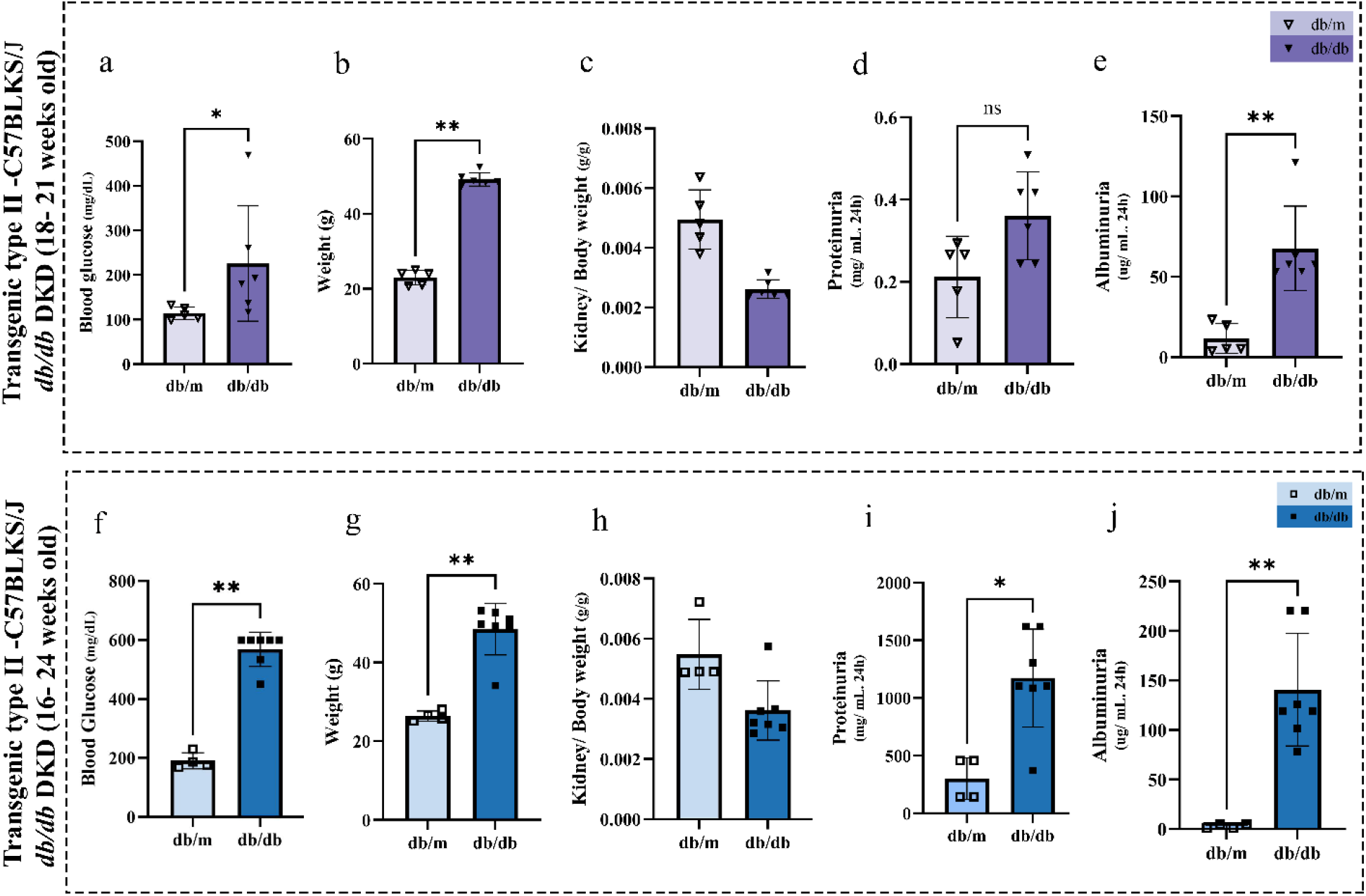
Biochemical characterization of the transgenic type 2 (C57BLKS/J) db/db DKD model across distinct age cohorts. Comparative metabolic profiles of early-stage (18–21 weeks old; n = 6 db/db, n = 5 db/m) and late-stage (16–24 weeks old; n = 7 db/db, n = 4 db/m) mice. (a, f) non-fasting blood glucose levels and (b, g) total body weight, both demonstrating highly significant increases in db/db mice across both age groups. (c, h) Kidney-to-body weight ratios, showing no statistically significant differences (ns) between diabetic and control groups. (d, i) 24-hour urinary total protein levels, showing a modest, non-significant trend in early-stage mice but a statistically significant elevation in the late-stage cohort. (e, j) 24-hour urinary albumin excretion, displaying 10-fold increase in the early-stage cohort and a more severe, >20-fold elevation in late-stage db/db mice. Data are shown as median with range. (ns = non-significant, * P < 0.05, **P < 0.01)

### 3.2 Comparative Histopathological Profiles in type 1 and type 2 mouse model

Renal histopathological alterations were evaluated across four parameters: glomerulomegaly, mesangial hypercellularity, tubular vacuolization, and arteriolar hyalinosis, and a total injury score was derived by aggregating these parameters within each cohort. The relative distribution and percentage of injury for markers for presence of lesion in DKD models were visualized using bar plots, and their statistical significance across the cohorts was comprehensively determined via Fisher’s exact test (Fig. 4) .Differences in the total injury score across the cohorts were comprehensively evaluated using the Mann-Whitney U test. To enhance the likelihood of detecting chronic, progressive structural alterations, mice with prolonged diabetes duration were selected. Additionally, only male mice were included to minimize sex-related biological variability and ensure uniform metabolic and histopathological comparisons. Initial non-blinded evaluations were performed to identify consistent pathological features, followed by blinded scoring analysis across all cohorts (STZ, Akita and db/db) to ensure objectivity.

Histological assessment of STZ-induced diabetic mice demonstrated the presence of all four evaluated lesions-glomerulomegaly, mesangial hypercellularity, arteriolar hyalinosis, and tubular vacuolization-none of which were identified in control animals (Fig. 3a-b, 4a). One diabetic mouse lacked glomerulomegaly despite exhibiting the remaining three lesions. Based on the cumulative evaluation, the total injury score was significantly elevated in the STZ group compared with its respective controls (Fig. 4b).

**Figure 3:**
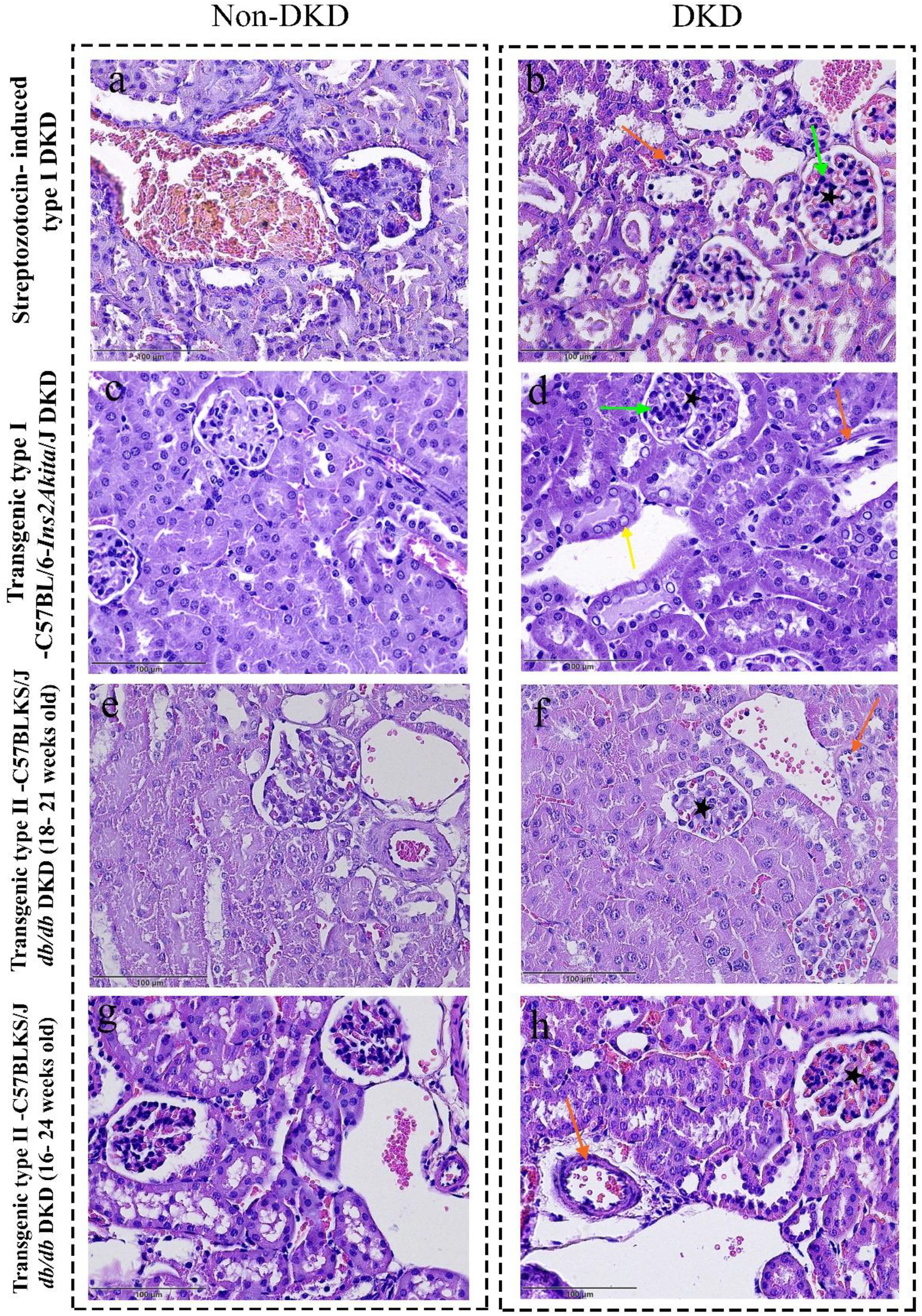
Comparative histopathological features across diabetic models. Representative H&E-stained kidney sections illustrate control (non-DKD; left column) and diabetic (DKD; right column) cohorts. (a, b) STZ-induced diabetic mice demonstrating diffuse renal injury, including glomerulomegaly, mesangial hypercellularity, arteriolar hyalinosis, which are absent in normal controls. (c, d) Akita mice showing glomerulomegaly, mesangial hypercellularity, arteriolar hyalinosis, and the presence of glycogenated nuclei, which was prominent in this model. (e, f) Early-stage db/db mice exhibiting mild and inconsistent histopathological changes. (g, h) Late-stage db/db mice showing variable renal lesions with less pronounced structural alterations compared with type 1 models. Pathological features are indicated as follows: glomerulomegaly (black stars), mesangial hypercellularity (green arrows), arteriolar hyalinosis (orange arrows), and glycogenated nuclei (yellow arrows).

**Figure 4:**
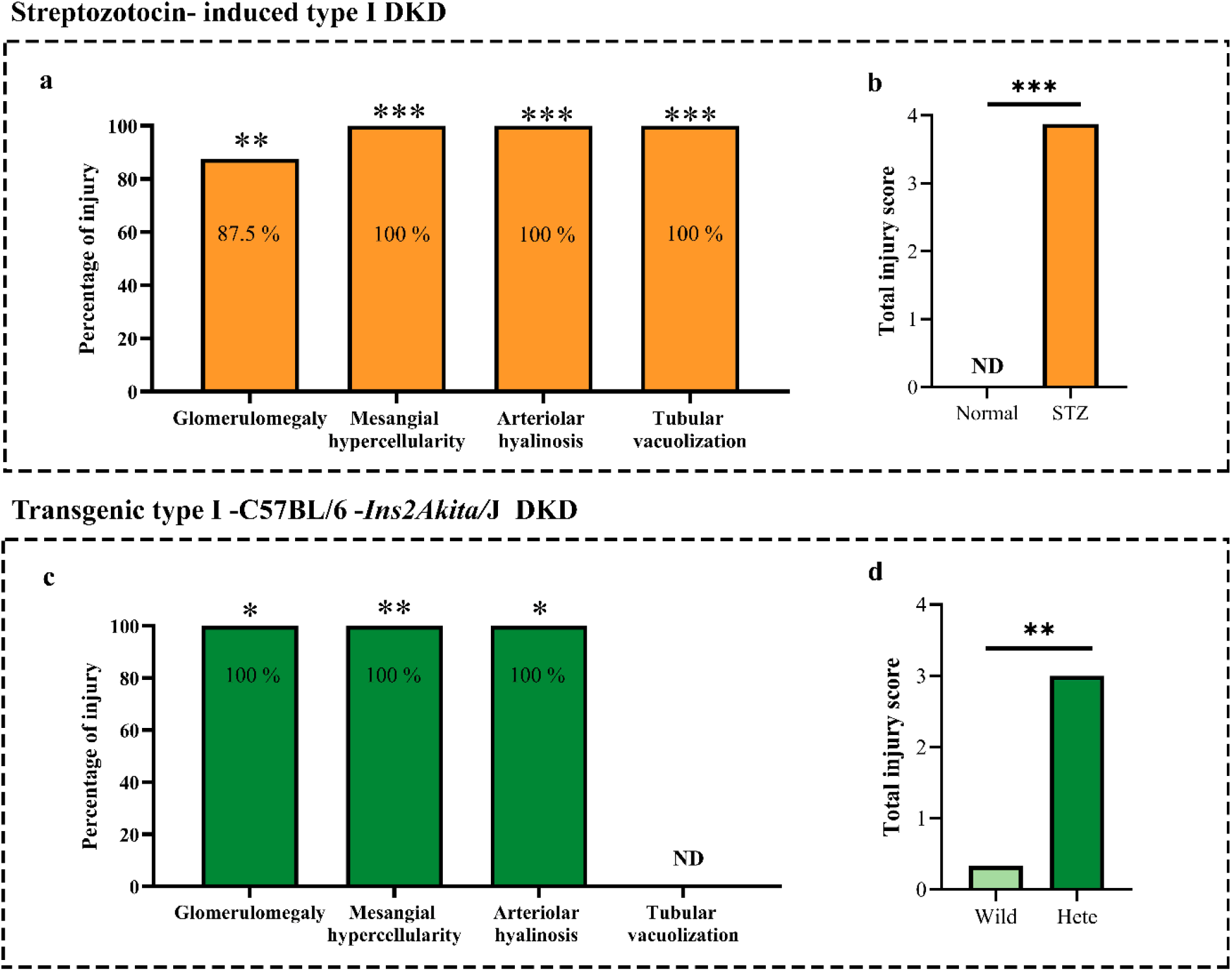
Categorical incidence and cumulative scoring of renal lesions across different diabetic mouse models. (a) Streptozotocin (STZ)-induced type 1 DKD model exhibiting a significant percentage of injury in all evaluated parameters, including tubular vacuolization. (b) Cumulative total injury score in the STZ model confirming a profound overall structural burden. (c) Transgenic type 1 (C57BL/6-Ins2Akita/J) DKD model showing prominent injury percentages in glomerulomegaly, mesangial hypercellularity, and arteriolar hyalinosis, while tubular vacuolization was not detected (ND). (d) Cumulative total injury score in the Akita (Hete) model demonstrating a significant overall histopathological alteration compared to wild-type controls. Histological evaluation was performed in a blinded manner, and data in (a, c) were analyzed using Fisher’s exact test, and data in (b, d) were analyzed via the Mann-Whitney U test. (* P < 0.05, ** P < 0.01, *** P < 0.001).

In the genetic type 1 Akita model, prominent structural damage was observed to the glomerular and vascular compartments, whereas tubular vacuolization was not detected (ND) (Fig. 3c-d, 4c). Control animals were largely lesion-free, though isolated glomerulomegaly and arteriolar hyalinosis were observed in two wild-type samples. The total injury score was significantly higher in Akita mice compared with wild-type controls (Fig. 4d). Additionally, glycogenated nuclei were detected within the renal parenchyma of Akita mice and, while also presenting the STZ model, were notably more prominent in this genetic model (Fig.3d).

Despite marked biochemical dysfunction, histopathological analysis of db/db mice revealed heterogeneous and inconsistent structural changes across the both age cohorts. In the early-stage cohort (18–21 weeks), glomerulomegaly, mesangial hypercellularity, and arteriolar hyalinosis were only variably detected without a consistent pattern of injury (Fig. 3e-f). Given the absence of uniform histopathological findings, a late-stage cohort was evaluated to determine whether structural lesions emerged with prolonged metabolic stress. However, despite more severe albuminuria and proteinuria in the late-stage db/db mice (Fig. 2i-j), histopathological findings remained similarly inconsistent, with no reliable increase in the prevalence of glomerulomegaly, mesangial hypercellularity, or arteriolar hyalinosis (Fig. 3g-h). Consequently, the statistical comparison of these histopathological parameter between the db/db and db/m mice remained highly inconsistent and non-significant in both age cohorts.

Given the discrepancy between the pronounced biochemical abnormalities and the relatively inconsistent light microscopic findings, ultrastructural analysis was performed to further characterize renal alterations in the early-stage db/db model (18-21 weeks old). Transmission electron microscopy revealed early glomerular ultrastructural changes, including irregular thickening of the glomerular basement membrane and focal podocyte foot process effacement compared with controls (Fig. 5a-d). These ultrastructural findings indicate that glomerular injury in early-stage db/db mice is present at the subcellular level prior to its detection by light microscopy, suggesting an early and pre-morphological stage of DKD progression.

**Figure 5:**
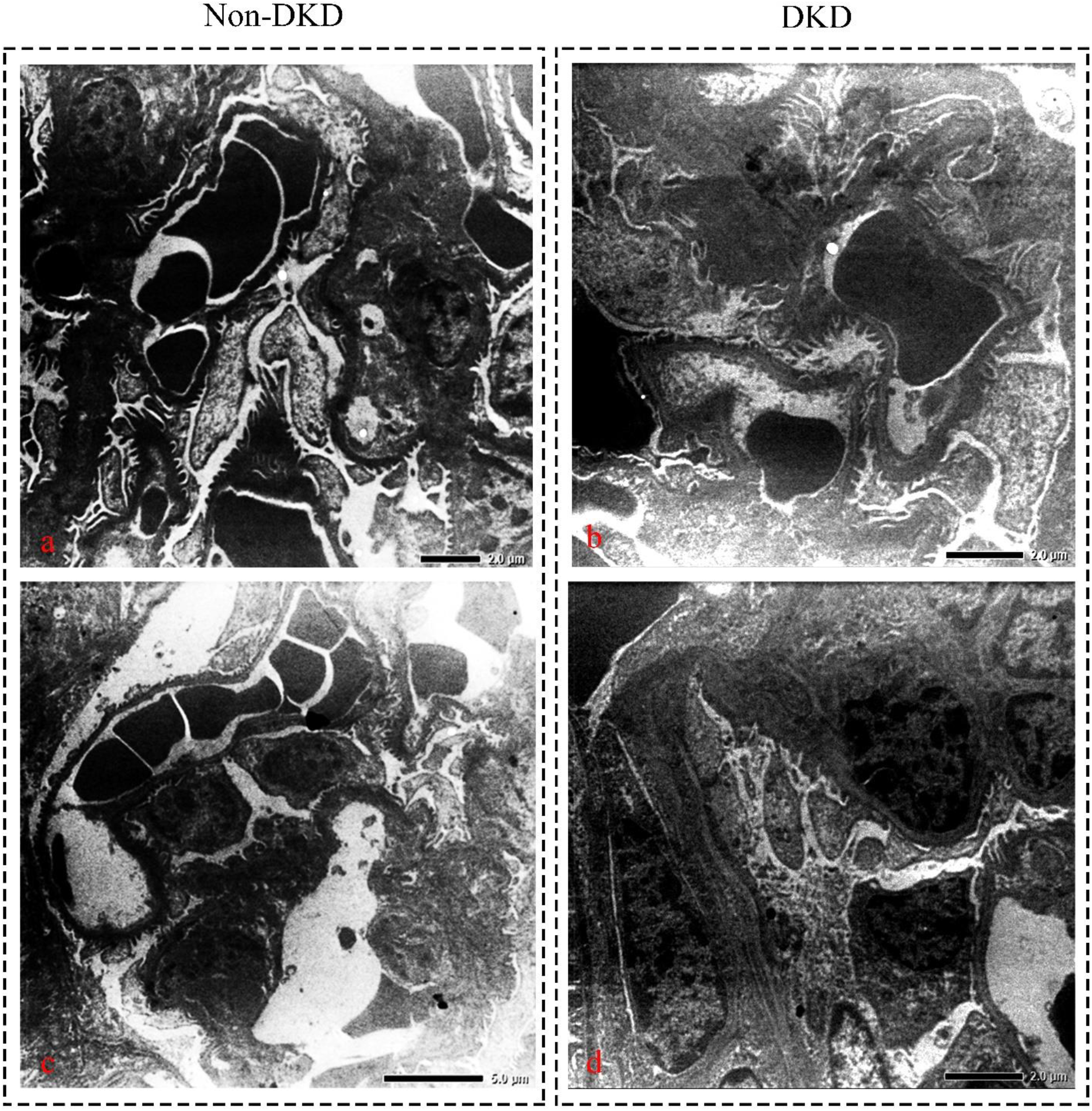
Transmission electron micrographs of db/db mouse glomerular ultrastructure. (a, c) Representative electron micrographs from control mice demonstrating intact glomerular ultrastructure, characterized by well-defined, distinct podocyte foot processes with preserved filtration slits and a uniform, thin glomerular basement membrane (GBM). (b, d) Micrographs from diabetic (db/db) mice revealing prominent pathological hallmarks of early-to-moderate DKD, including extensive, diffuse podocyte foot process effacement and irregular, pronounced thickening of the GBM. Scale bars are indicated individually in each panel (2.0 µm for a,b,d and 5.0 µm for c).

## 4. Discussion

Diabetic kidney disease (DKD) represents one of the most heterogeneous complications of diabetes mellitus, and no single animal model fully recapitulates the complete spectrum of human pathology [5]. Previous studies have emphasized that available rodent models reproduce only selective features of DKD, with phenotypic expression varying considerably according to genetic background, induction method, and disease duration [7, 16]. While extensive literature focuses on advanced structural hallmarks like nodular glomerulosclerosis and diffuse tubulointerstitial fibrosis and etc., these features were completely absent under routine light microscopy in our baseline cohorts. Instead, the novel strength of this study lies in capturing alternative, early-to-moderate histopathological changes that are frequently overlooked or less routinely quantified in DKD animal experiments. By focusing on the categorical of some overlooked specific indices like glomerulomegaly, mesangial hypercellularity, arteriolar hyalinosis, and tubular vacuolization, our findings demonstrate specific structural phenotypes despite comparable downstream metabolic and functional disturbances across type 1 (STZ-induced and Akita) and type 2 (db/db) DKD animal model. The alternative parameters evaluated herein provided a reliable, multi-compartmental assessment of renal injury. Glomerulomegaly reflects early glomerular hypertrophy, while mesangial hypercellularity serves as a clear indicator of initial mesangial expansion and glomerular tuft stress [17]. Tubular vacuolization serves as a marker of tubular epithelial damage and metabolic overload [18], while arteriolar hyalinosis reflects chronic microvascular damage within the renal vasculature [19].

Both the STZ and Akita models showed consistent structural alterations in the present study. the STZ model exhibited all four evaluated lesions **-** namely glomerulomegaly, mesangial hypercellularity, arteriolar hyalinosis, and tubular vacuolization-indicating a more diffuse pattern of renal injury. This finding is consistent with previous studies showing that STZ-induced diabetes can lead to glomerular hypertrophy, mesangial expansion, arteriolar hyalinosis, and tubular damage, depending on strain susceptibility and experimental conditions [20]. Collectively, these findings suggest that while both models effectively reproduce structural changes associated with type 1 diabetes, the STZ model more comprehensively captures the full spectrum of histopathological changes assessed in this study.

In contrast to STZ-induced mice, Akita mice, glomerulomegaly, mesangial hypercellularity, and arteriolar hyalinosis were observed, whereas tubular vacuolization was notably absent. As shown, glomerular and vascular lesions were prominent in the Akita model, which is consistent with earlier reports. However, these reports describe additional markers that were not detected in this study, such as mesangial expansion, glomerular basement membrane (GBM) thickening, and progressive glomerular changes in this model [21]. As previously noted, advanced lesions like widespread glomerulosclerosis are usually restricted in Akita mice, suggesting that this model represents moderate rather than severe DKD progression [7]. Furthermore, glycogenated nuclei were detected in both the Akita and STZ mouse models, though they were qualitatively more pronounced in the Akita mice. Intranuclear accumulation of glycogen aggregates within proximal tubule cells-an indicator of tubular damage-has been reported in STZ-induced models, however, this finding has not been previously described in Akita mice [22]. The localized presence of glycogenated nuclei in renal epithelial cells is a unique morphological indicator of severe, sustained intracellular metabolic stress, reflecting advanced metabolic injury under chronic hyperglycemic states [23, 24].

Compared to type 1 diabetes models, the db/db model demonstrated a markedly distinct pattern. Despite pronounced metabolic abnormalities, including obesity, hyperglycemia, and albuminuria, histopathological changes assessed by light microscopy were heterogeneous and inconsistent across both the early-stage (18–21 weeks old) and late-stage (16–24 weeks old) cohorts.. The inclusion of two age groups was proposed to evaluate whether disease progression would enhance histopathological detectability; however, variability persisted even in late-stage animals.

Notably, the observed discrepancy between robust biochemical alterations and limited histological changes in db/db mice highlights a known limitation of experimental DKD models. As reported in previous studies, many genetic models of both type 1 and type 2 diabetes predominantly reflect early-stage disease and may fail to develop overt structural lesions without additional modifiers such as aging, hypertension, or secondary renal stressors [7, 25, 26]. In this context, our ultrastructural analysis provides additional insight. Transmission electron microscopy revealed early glomerular alterations, including thickening of GBM and focal effacement of podocyte foot process. These are well-established ultrastructural features of early DKD that may not be detectable by routine light microscopy.

This notable divergence between our findings and previous reports can be primarily attributed to inherent biological differences across animal strains, variations in experimental conditions, disease duration, and the age at which renal tissues were evaluated. The conventional histopathological parameters in the literature, which typically require prolonged disease duration or accelerated disease progression to manifest consistently, the indices in our study exhibited highly consistent phenotypes in type 1 experimental animal model. Consequently, the alternative parameters identified-glomerulomegaly, mesangial hypercellularity, arteriolar hyalinosis, and tubular vacuolization-appear to be more sensitive and reliable indicators of early-to-moderate DKD progression. These features can be readily identified by routine light microscopy, even when classical late-stage fibrotic markers, such as tubulointerstitial fibrosis and glomerulosclerosis, are not prominently developed because of model-specific differences.

## 5. Conclusion

In summary, diabetic mouse models exhibit substantial differences in their ability to reproduce structural features of human DKD. The alternative histopathological indices evaluated in this study-namely glomerulomegaly, mesangial hypercellularity, tubular vacuolization, and arteriolar hyalinosis-proved highly capable of delineating mild-to-moderate tissue injury, offering a practical framework for future experimental validation. These results highlight that no single model fully captures the complexity and full spectrum of DKD. Therefore, appropriate model selection is essential for a more accurate and comprehensive evaluation of renal injury in experimental diabetes.

## Statements & Declarations

### Ethics approval

All animal experiments were conducted in accordance with the NIH Guide for the Care and Use of Laboratory Animals and approved by the relevant institutional ethics committees prior to study initiation (University of Nizwa Ethics Committee: VCGSR, AREC/01/202 for Akita and db/db models; Isfahan University of Medical Sciences Ethics Committee: IR.MUI.AEC.1401.046 for STZ-induced models.

### Author contributions

R.R. (Raziyeh Rezaei): Methodology, Formal analysis, Investigation, Data curation, Writing - Original Draft, Writing - Review & Editing and Funding acquisition.

A.N. (Azar Naiemi): Conceptualization, Investigation, Data curation, Writing - Review & Editing, Validation.

Y.G. (Yousof Gheisari): Conceptualization, Supervision, Methodology, Validation, Resources, Data curation, Writing - Original Draft, Writing - Review & Editing, Visualization, Project administration.

Z.R. (Zahra Ramazani): Methodology, Formal analysis, Investigation, Data curation, Writing - Original Draft, Writing - Review & Editing.

F.J.-A. (Fatemeh Jamshidi-Adegani): Methodology, Investigation, Data curation, Writing - Original Draft, Writing - Review & Editing and Funding acquisition.

I.S.A.-A. (Issa Sulaiman Al-Amri): Investigation, Methodology, Writing - Review & Editing.

H.D. (Hoda Doustmohammadi): Investigation, Methodology, Writing - Review & Editing.

S.A.-H. (Sulaiman Al-Hashmi): Conceptualization, Supervision, Methodology, Validation, Resources, Data curation, Writing - Original Draft, Writing - Review & Editing, Visualization, Project administration.

## Acknowledgements

Not applicable

## References

1. Ortiz, A., et al., The updated global burden of chronic kidney disease: one death every 20 seconds. Nephrology Dialysis Transplantation, 2026.

2. Qasim, H., et al., Histopathology of Diabetic Nephropathy: Beyond Glomerular Basement Membrane Thickening. Cureus, 2025. 17(9): p. e93497.

3. Espinel, E., et al., Renal Biopsy in Type 2 Diabetic Patients. J Clin Med, 2015. 4(5): p. 998–1009.

4. Breyer, M.D., et al., Mouse models of diabetic nephropathy. J Am Soc Nephrol, 2005. 16(1): p. 27–45.

5. Xie, K., et al., Research and advances in mouse models of diabetic nephropathy: a narrative review. BMC Nephrol, 2025. 26(1): p. 511.

6. Wu, J. and L.J. Yan, Streptozotocin-induced type 1 diabetes in rodents as a model for studying mitochondrial mechanisms of diabetic β cell glucotoxicity. Diabetes Metab Syndr Obes, 2015. 8: p. 181–8.

7. Kitada, M., Y. Ogura, and D. Koya, Rodent models of diabetic nephropathy: their utility and limitations. Int J Nephrol Renovasc Dis, 2016. 9: p. 279–290.

8. Tesch, G.H. and T.J. Allen, Rodent models of streptozotocin-induced diabetic nephropathy (Methods in Renal Research). Nephrology, 2007. 12(3): p. 261–266.

9. Chang, J.H. and S.B. Gurley, Assessment of diabetic nephropathy in the Akita mouse. Methods Mol Biol, 2012. 933: p. 17–29.

10. Wang, B., P.C. Chandrasekera, and J.J. Pippin, Leptin- and leptin receptor-deficient rodent models: relevance for human type 2 diabetes. Curr Diabetes Rev, 2014. 10(2): p. 131–45.

11. Giralt-López, A., et al., Revisiting Experimental Models of Diabetic Nephropathy. Int J Mol Sci, 2020. 21(10).

12. National Research Council. Guide for the Care and Use of Laboratort Animals. 8th ed. Washington (DC): National Academies Press (US). 2011. doi: 10.17226/12910.

13. Furman, B.L., Streptozotocin-Induced Diabetic Models in Mice and Rats. Curr Protoc, 2021. 1(4): p. e78.

14. Barber, A.J., et al., The Ins2Akita mouse as a model of early retinal complications in diabetes. Invest Ophthalmol Vis Sci, 2005. 46(6): p. 2210–8.

15. Prakash, M., et al., Determination of urinary peptides in patients with proteinuria. Indian J Nephrol, 2008. 18(4): p. 150–4.

16. Bufi, R. and R. Korstanje, The impact of genetic background on mouse models of kidney disease. Kidney Int, 2022. 102(1): p. 38–44.

17. Kataoka, H., K. Nitta, and J. Hoshino, Glomerular hyperfiltration and hypertrophy: an evaluation of maximum values in pathological indicators to discriminate “diseased” from “normal”. Front Med (Lausanne), 2023. 10: p. 1179834.

18. Zhou, C., A.J. Yool, and R.W. Byard, Basal Vacuolization in Renal Tubular Epithelial Cells at Autopsy and Their Relation to Ketoacidosis. Journal of Forensic Sciences, 2017. 62(3): p. 681–685.

19. Menon, R., et al., Defining the molecular correlate of arteriolar hyalinosis in kidney disease progression by integration of single cell transcriptomic analysis and pathology scoring. medRxiv, 2023: p. 2023.06.14.23291150.

20. Luo, W., et al., Translation Animal Models of Diabetic Kidney Disease: Biochemical and Histological Phenotypes, Advantages and Limitations. Diabetes Metab Syndr Obes, 2023. 16: p. 1297–1321.

21. Itano, S., et al., Non-purine selective xanthine oxidase inhibitor ameliorates glomerular endothelial injury in Ins(Akita) diabetic mice. Am J Physiol Renal Physiol, 2020. 319(5): p. F765–f772.

22. Glastras, S.J., et al., Mouse Models of Diabetes, Obesity and Related Kidney Disease. PLOS ONE, 2016. 11(8): p. e0162131.

23. Kang, J., et al., Glycogen accumulation in renal tubules, a key morphological change in the diabetic rat kidney. Acta Diabetol, 2005. 42(2): p. 110–6.

24. Sullivan, M.A. and J.M. Forbes, Glucose and glycogen in the diabetic kidney: Heroes or villains? EBioMedicine, 2019. 47: p. 590–597.

25. Zeng, M., et al., Animal models for diabetic kidney disease: perspectives and prospects. Frontiers in Veterinary Science, 2026. Volume 13-2026.

26. Li, F., et al., Optimizing diabetic kidney disease animal models: Insights from a meta-analytic approach. Animal Model Exp Med, 2023. 6(5): p. 433–451.

